# Structural characterization of HIV-1 matrix mutants implicated in envelope incorporation

**DOI:** 10.1101/2020.10.14.339457

**Authors:** Gunnar N. Eastep, Ruba H. Ghanam, Todd J. Green, Jamil S. Saad

**Affiliations:** Department of Microbiology, University of Alabama at Birmingham, Birmingham, AL 35294

**Author notes:** To whom correspondence should be addressed: Jamil S. Saad, Ph.D., 845 19^th^ Street South, Birmingham, AL 35294.

**Keywords:** Human immunodeficiency virus type 1 (HIV-1), retroviruses, virus assembly, Gag polyprotein, myristoylated matrix (MA), plasma membrane (PM), phosphatidylinositol 4,5-bisphosphate (PI(4,5)P_2_), Envelope protein (Env), gp41 cytoplasmic tail (gp41CT), nuclear magnetic resonance (NMR), x-ray crystallography, analytical ultracentrifugation (AUC)

## Abstract

During the late phase of HIV-1 infection, viral Gag polyproteins are targeted to the plasma membrane (PM) for assembly. Gag localization at the PM is a prerequisite for the incorporation of the envelope protein (Env) into budding particles. Gag assembly and Env incorporation are mediated by the N-terminal myristoylated matrix (MA) domain of Gag. Nonconservative mutations in the trimer interface of MA (A45E, T70R, and L75G) were found to impair Env incorporation and infectivity, leading to the hypothesis that MA trimerization is an obligatory step for Env incorporation. Conversely, Env incorporation can be rescued by a compensatory mutation in the MA trimer interface (Q63R). The impact of these MA mutations on the structure and trimerization properties of MA is not known. In this study, we employed NMR spectroscopy, x-ray crystallography, and sedimentation techniques to characterize the structure and trimerization properties of HIV-1 MA A45E, Q63R, T70R, and L75G mutant proteins. NMR data revealed that these point mutations did not alter the overall structure and folding of MA but caused minor structural perturbations in the trimer interface. Analytical ultracentrifugation data indicated that mutations had a minimal effect on the MA monomer–trimer equilibrium. The high-resolution x-ray structure of the unmyristoylated MA Q63R protein revealed hydrogen bonding between the side chains of Arg-63 and Ser-67 located in the center of the trimer interface, providing the first structural evidence for a stabilization of the trimer form. These findings advance our knowledge of the interplay of MA trimerization and Env incorporation into HIV-1 particles.

---

During the late phase of HIV-1 replication, the viral Gag polyproteins are targeted to the plasma membrane (PM) for particle budding and release (1–7). Binding of HIV-1 Gag polyproteins to the inner leaflet of the PM is mediated by the matrix (MA) domain, which harbors a bipartite signal represented by an N-terminal myristoyl group (myr) and a highly basic region (HBR). An abundance of studies demonstrated that HIV-1 Gag association with membranes is regulated by multiple factors including electrostatic and hydrophobic interactions, protein multimerization, cellular and viral RNA, specific phospholipids such as phosphatidylinositol 4,5-bisphosphate (PI(4,5)P_2_) (7–13). Gag binding to membranes is also enhanced by inclusion of phosphatidylserine (PS) and cholesterol (6,8,9,12,14–20).

Pioneering x-ray crystallography studies of HIV-1 unmyristoylated MA [myr(–)MA] revealed that the protein adopts a trimer structure (21). However, solution NMR studies indicate that the myr(–)MA protein is monomeric at all tested concentrations (22–27). On the other hand, sedimentation equilibrium data confirmed that the MA protein resides in monomer–trimer equilibrium. NMR–based structural studies have shown that the myr group of MA can adopt sequestered and exposed conformations (26). Extrusion of the myr group can be increased by factors that promote protein self-association, such as increasing protein concentration or inclusion of the capsid (CA) domain (26). Subsequent studies revealed that myr exposure in MA is promoted by binding of PI(4,5)P_2_ (25) and is regulated by pH (27). In a recent study, we engineered a stable recombinant HIV-1 MA trimer construct by fusing a foldon domain (FD) of phage T4 fibritin to the MA C terminus (28). Hydrogen–deuterium exchange MS data supported a MA–MA interface that is consistent with that observed in the crystal structure of the myr(–)MA trimer (28). A plethora of structural and biophysical studies have provided invaluable insights into retroviral MA proteins binding to phospholipids and membrane mimetics (23–25, 29–38). These studies identified key molecular determinants of MA–membrane interactions and advanced our knowledge of retroviral assembly.

Studies by Barklis and co-workers revealed that the HIV-1 MA and MACA proteins assemble as hexagonal cage lattices on PS/cholesterol membrane monolayers and as hexamers of trimers in the presence of PI(4,5)P_2_ (39–41). The hexamer of trimers assembly is currently used as a model to explain the mechanism by which the envelope (Env) protein is incorporated into virus particles (42–45). In its processed form, Env consists of two non-covalently associated subunits, a surface glycoprotein (gp120) and a transmembrane domain (gp41), which are products of proteolytic cleavage of the viral gp160 precursor protein (reviewed in (46–48)). There is strong evidence that the cytoplasmic tail of gp41 (gp41CT) plays a functional role in Env incorporation in physiologically relevant cell types (42,44,45,49–52). NMR studies of HIV-1 gp41CT revealed that the N-terminal 45 residues lack secondary structure and are not associated with the membrane. The C-terminal 105 residues, however, form three membrane–bound amphipathic α-helices with distinctive structural features such as variable degree of membrane penetration, hydrophobic and basic surfaces, clusters of aromatic residues, and a network of cation–*π* interactions (53).

Genetic and biochemical studies suggested that the MA domain of Gag and the gp41CT play distinct, yet complementary, roles in Env incorporation into budding particles (42,44,45,49,50,54–59). Point mutations in MA (L13E, E17K, L31E, V35E, or E99V) were found to impair HIV-1 Env incorporation (42–44, 50, 56). [Note: The N-terminal Met, which is absent in the myristoylated protein, is designated as residue 1. In contrast, other studies considered the N-terminal Gly of the myristoylated protein as residue 1]. When mapped on the putative hexamer of trimer model of MA, these residues were found to point toward the hexamer centers (Fig. 1) (39, 40). It was also shown that substitution of residue Gln^63^ with Arg suppressed Env incorporation defects of the L13E, E17K, L31E, V35E, or E99V MA mutations and of a gp41CT mutation that has the same phenotype (42–44, 50). Freed and co-workers provided biochemical evidence that MA trimerization is an obligatory step in the assembly of infectious HIV-1 and demonstrated a correlation between loss of MA trimerization and loss of Env incorporation (44). It was suggested that Q63R mutation may stabilize the trimer structure such that MA lattices, which form large hexamer holes, are favored over those that feature small hexamer holes (Fig. 1) (44). In another study, biochemical and pulldown experiments suggested that Q63R mutation promoted interaction with gp41CT, and this mutation did not alter the organization of MA on a membrane layer (41). Non-conservative mutations at positions Ala^45^, Ser^72^, and Leu^75^ located in the trimer interface (Fig. 1) also impaired Env incorporation and infectivity, and biochemical cross-linking experiments showed that these mutations reduced trimerization of MA (44). Although genetic and biochemical studies provided evidence for a correlation between Env incorporation and formation of MA trimers, the structural or biophysical evidence for such a correlation is lacking. In particular, it is not known how single residue mutations which impair or rescue Env incorporation impact the structure and oligomerization properties of MA.

**Figure 1.**
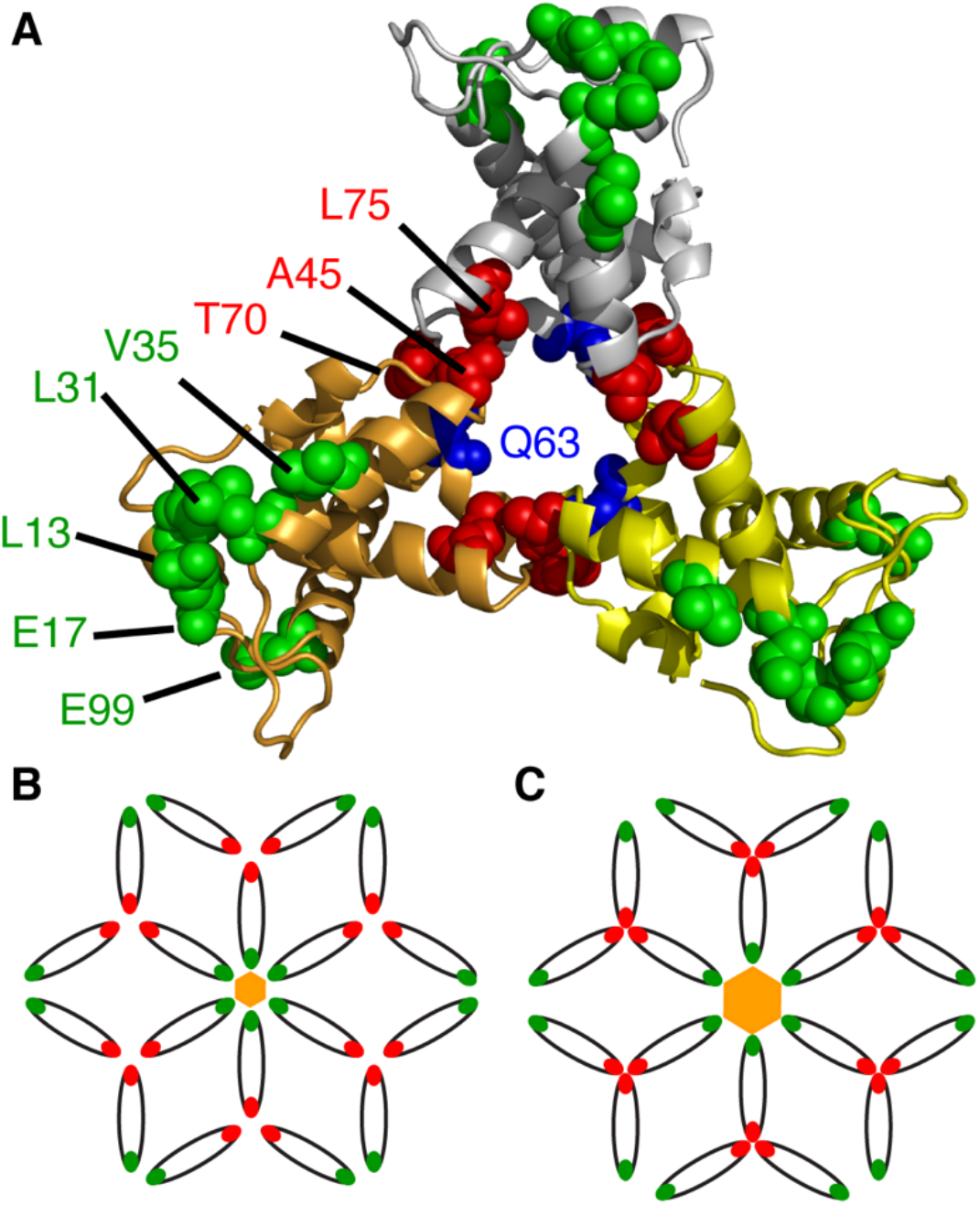
MA trimer model and residues implicated in Env incorporation. (A) Structure of the myr(–)MA trimer (PDB ID: 1HIW) showing residues implicated in Env incorporation. Gln^63^ is shown as blue sticks. (B) Schematic representation of the hexamer of trimer arrangement of MA based on studies by Alfadhli *et al*. (39) and adopted by Freed *et al*. (44). In this model, a ~45 nm central aperture is formed by residues at the tip of the hexamer (green). It is suggested that the gp41CT protein is accommodated in the central aperture. Mutations of residues in the trimer interface (red sticks) significantly disrupted Env incorporation. (C) Perturbations of the putative hexamer or trimer interface in the MA lattice are thought to create smaller central aperture (~30 nm) that may cause a steric exclusion of gp41CT.

In this report, we employed NMR spectroscopy, x-ray crystallography and analytical ultracentrifugation (AUC) techniques to characterize the structure and trimerization properties of HIV-1 MA mutants that have been shown to impair Env incorporation (L75G, A45E, and T70R) or suppress Env incorporation defects (Q63R). We show that these mutations had no adverse effect on the structure or folding of MA and had only a minimal effect on the monomer– trimer equilibrium. NMR data revealed that these mutations caused chemical shift perturbations (CSPs) for residues located in the trimer interface. The x-ray structure of myr(–)MA Q63R protein revealed hydrogen bonding between the side chains of Arg^63^ and Ser^67^, which are located in the trimer interface. These findings advance our knowledge of the interplay of MA trimerization and Env incorporation into HIV-1 particles.

## Results

### Effect of point mutations on the structure of MA

Point mutations in the HIV-1 MA protein (A45E, Q63R, T70R and L75G) were made by site-directed mutagenesis of the MA gene embedded in a co-expression vector harbouring yeast N-myristoyltransferase (26). Point mutations had no detectable effect on protein expression or the efficiency of myristoylation. Two-dimensional (2D) ^1^H-^15^N HSQC NMR spectrum provides a structure-sensitive signal for each amide group in a protein, and thus, it can be used to report on changes to protein structure caused by amino acid substitutions. As shown in Figures 1 and S1, the HSQC spectra for the four MA mutants are very similar to the spectrum of the wild-type (WT) protein, indicating that amino acid substitutions had no adverse effect on the structure and folding of the protein. This result was supported by the ^1^H, ^15^N, Cα, and Cβ chemical shifts, which were very similar to the corresponding shifts of the WT MA protein (25, 26).

### Effect of amino acid substitutions on the myr switch

Previous NMR studies of the WT MA protein have shown that a subset of ^1^H and ^15^N resonances for amino acid residues 2-18, Val^35^, Trp^36^, Arg^39^, Gly^49^, Glu^52^, and His^89^ shift progressively toward the corresponding frequencies observed for myr(–)MA upon increasing protein concentration. These shifts were attributed to exposure of the myr group and a concomitant shift in the monomer–trimer equilibrium toward the trimeric species (26). Subsequent NMR studies rvealed that point mutations in the N-terminal region of MA (e.g., V7R, L8A or L8I) that impaired membrane targeting of Gag and inhibited virus assembly and release (60–63), did not exhibit concentrationdependent myristate exposure (24). Structural data revealed conformational changes that appeared to be responsible for stabilizing the myristate–sequestered species and inhibiting exposure (24). Herein, we assessed whether the A45E, Q63R, T70R and L75G mutations in MA had any effect on the positioning of the myr group and/or the concentration–dependent myr-exposure. We obtained ^1^H-^15^N HSQC spectra for WT and A45E, Q63R, T70R, and L75G mutant MA proteins at three protein concentrations (50, 150 and 450 μM) (Fig. 2). Like WT MA, for all mutants the ^1^H and ^15^N resonances for residues 2-18, Val^35^, Trp^36^, Arg^39^, Gly^49^, Glu^52^, and His^89^ were sensitive to protein concentration and shifted progressively toward the corresponding frequencies observed for myr(–)MA upon increasing protein concentration (Fig. 2). This result indicates that none of the four amino acid substitutions had any detectable effect on the concentration–dependent myr switch.

**Figure 2.**
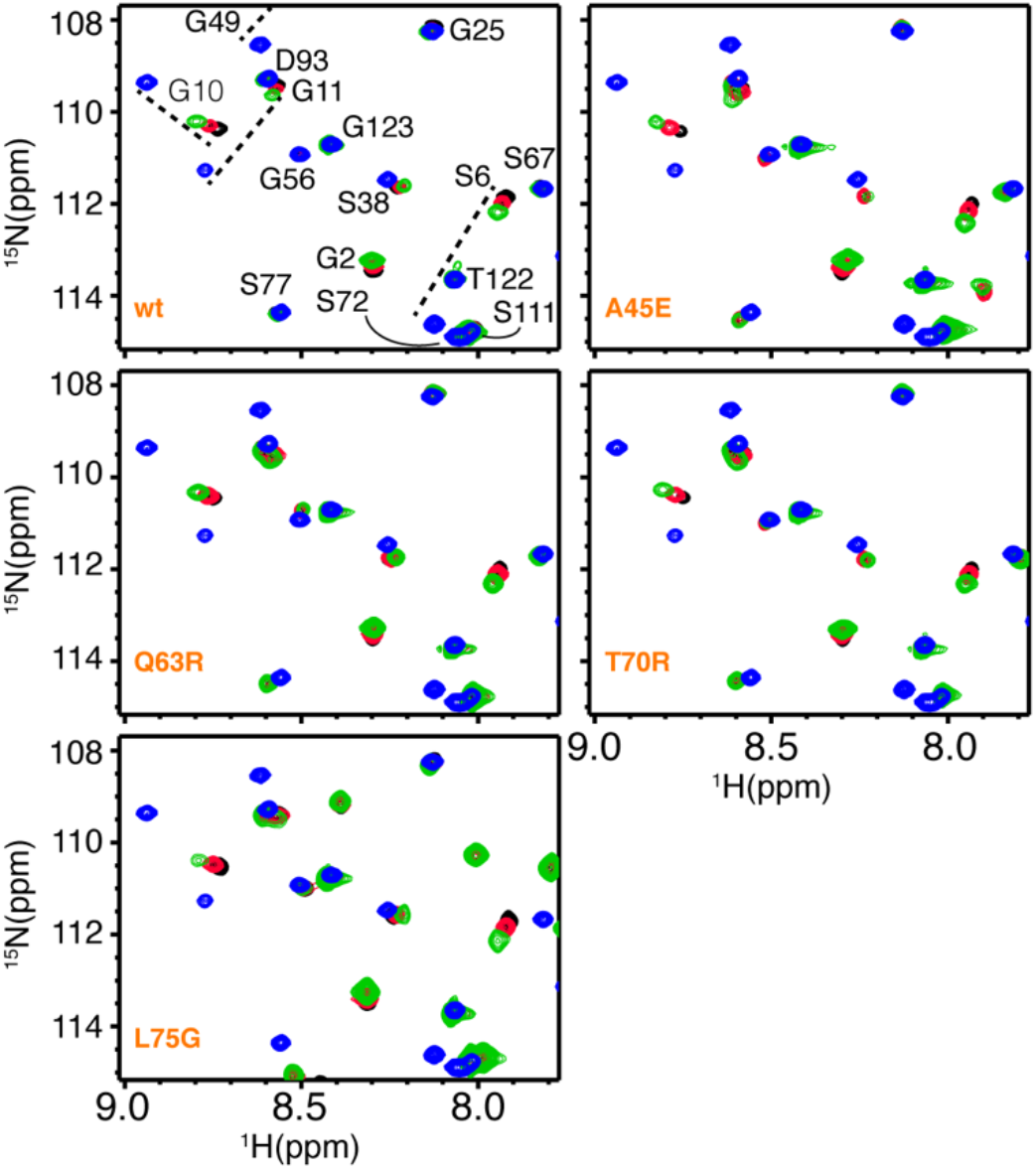
NMR spectra of WT and mutant MA at different protein concentrations. Overlay of 2D ^1^H-^15^N HSQC spectra for WT, myr(–), and mutant MA proteins collected at different concentrations [50 μM (black), 150 μM (red), and 450 μM (green)]. Like WT MA, for all four mutant proteins a subset of ^1^H-^15^N resonances of MA shifted toward the corresponding signals of myr(–)MA (blue), indicating a shift in equilibrium towards the myr–exposed state.

### Chemical shift comparison between the WT and mutant MA proteins

We mapped the chemical shift changes caused by amino acid substitutions in MA and assessed whether they induced structural and/or conformational changes in distal regions of the MA protein, especially within the trimer interface. Typically, only a few signals corresponding to amino acids in the vicinity of the mutation site exhibit major chemical shift changes in the ^1^H-^15^N HSQC spectrum. Chemical shift differences between the WT and mutant MA proteins is provided in Figure 3, and a cartoon representation of the X-ray structure of the WT MA protein onto which the locations of residues that exhibited significant amide ^1^H-^15^N chemical shift change are mapped (Fig. 4). As expected, the largest differences are observed in the vicinity of mutation site. Interestingly, other regions that are distant from the mutation site were also perturbed. For the A45E mutant, in addition to the CSPs observed for signals of residues 39-47 which are in the vicinity of the mutation site, resonances corresponding to Glu^74^ and Leu^75^ also exhibited significant CSPs (Fig. 3). Interestingly, these two residues are quite distant from the mutation site within the same molecule but are intermolecularly adjacent (~4.5 Å) in the trimer structure of myr(–)MA (Fig. 4). This result suggests that substitution of Ala^45^ to Glu may have caused structural perturbations in the trimer interface. Next, we examined the Q63R mutant. Gln^63^ is located in helix 4, with its side chain projecting towards the loop comprised of residues 42-46 (~4.1 Å to residue Val^46^) within the same molecule. As shown in Figure 3, in addition to the mutation site two other regions exhibited significant CSPs. These comprise residues 42-47 and 74-76. Mapping out the CSPs on the trimer structure of myr(–)MA revealed that this mutation can cause structural perturbations for residues 42-46 within the same molecule (Fig. 4). Therefore, it is likely that the CSPs corresponding to residues 42-46 are caused by intramolecular structural adjustments or perturbations as a result of this substitution. However, whereas amino acid residues 74-75 are intramolecularly distant from Gln^63^, these residues are located in the trimer interface (Fig. 4). In this case, the effect of Q63R mutation could be direct on residues 74-76 or indirect through a local structural perturbation of residues 42-47, which are also close to residues 74-76 (Fig. 4).

**Figure 3.**
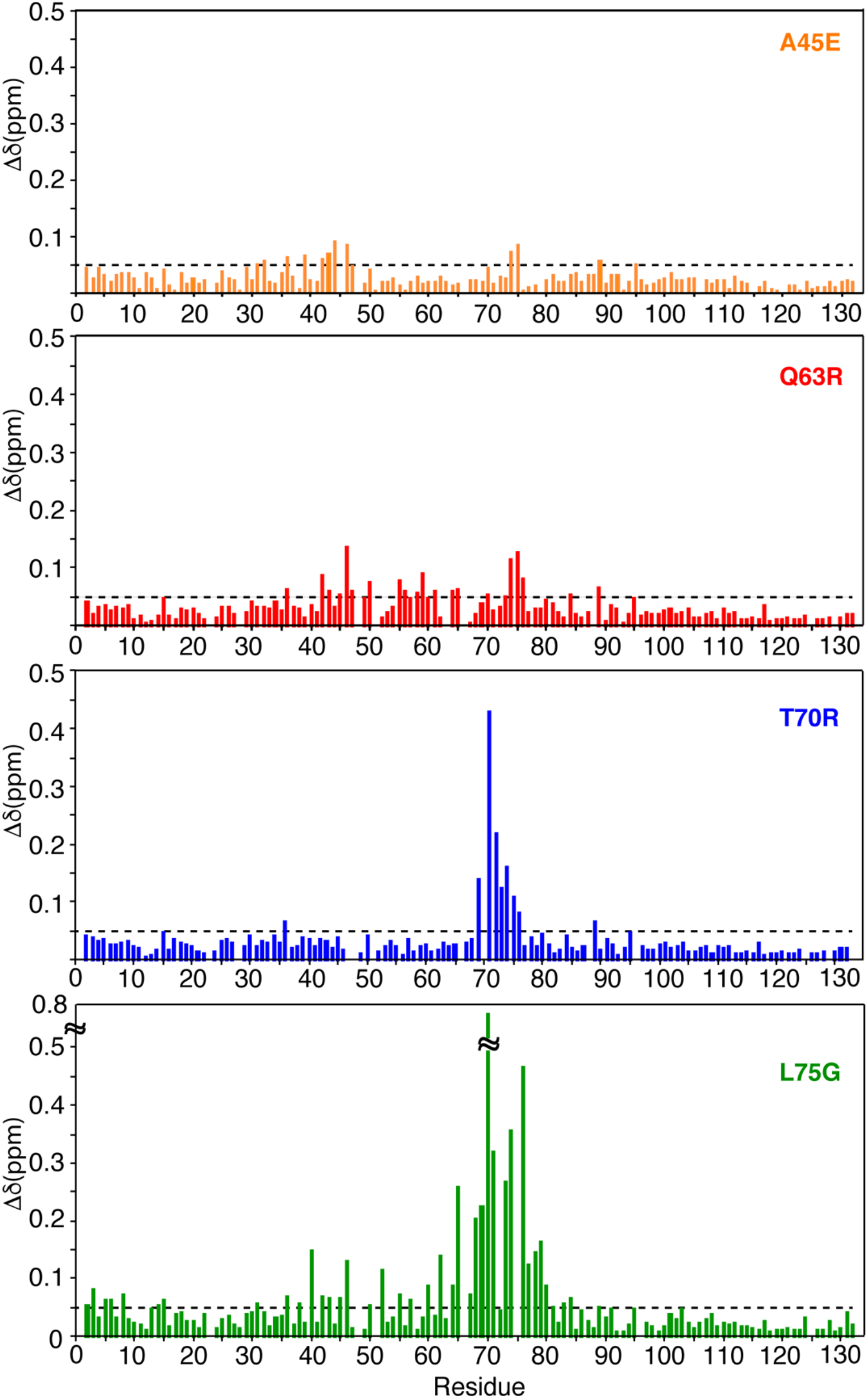
Chemical shift differences between the WT and mutant MA proteins. Normalized ^1^H-^15^N chemical shift differences are plotted vs. residue number. Δδ > 0.05 ppm indicates significant chemical shift changes.

**Figure 4.**
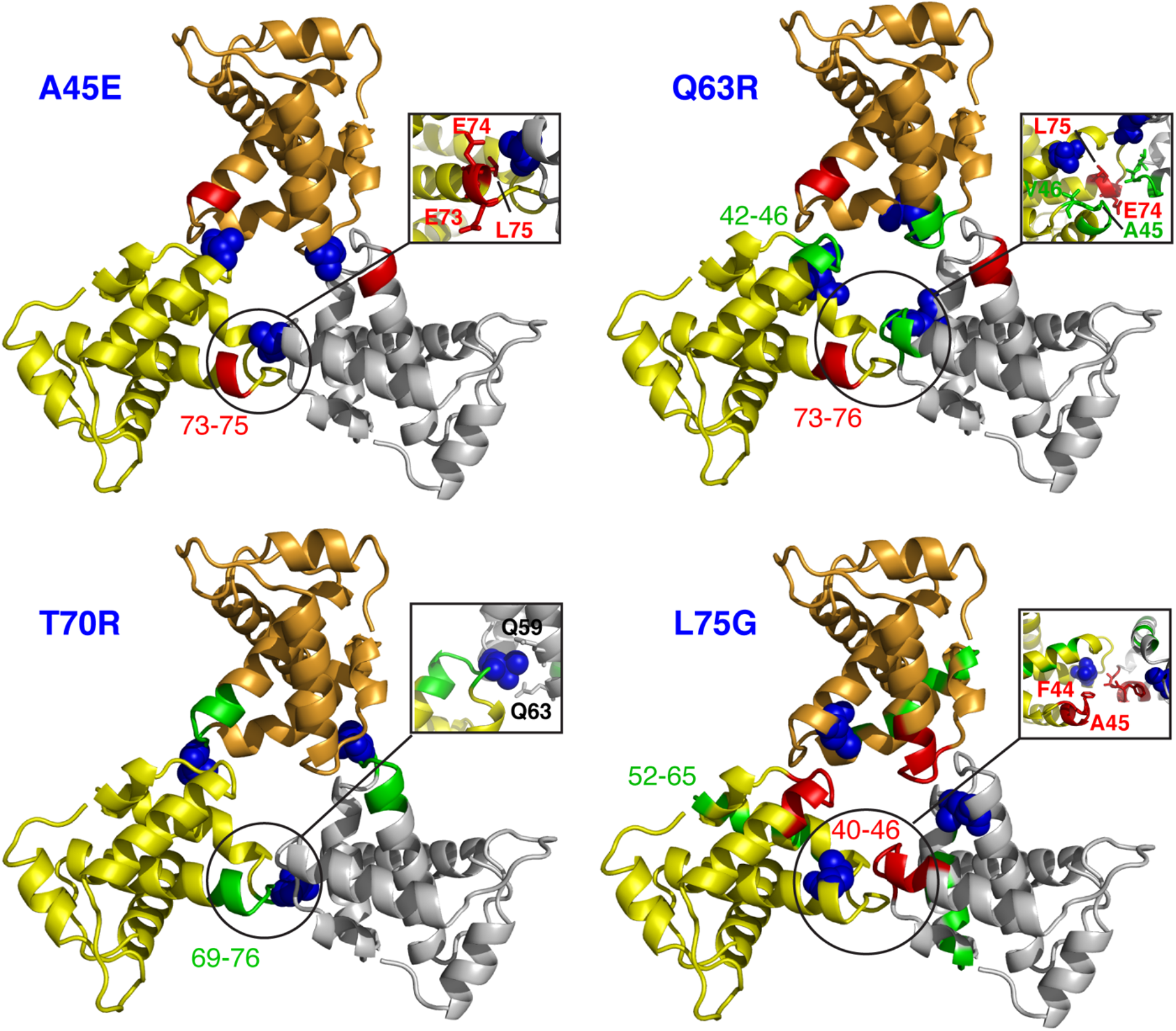
Chemical shift mapping on the structure of the myr(–)MA trimer. Structural mapping of the chemical shift differences onto the X-ray structure of WT MA trimer (PDB ID: 1HIW). The location of the amino acid mutation is marked with a blue sphere. Positions of residues whose amide resonances exhibited Δδ > 0.05 ppm are highlighted in red (intermolecular) and green (intramolecular).

Among the four MA mutants, the largest CSPs have been observed in the HSQC spectra of the T70R and L75G proteins (Fig. 3). For T70R, the most significant CSPs are for residues localized in the vicinity of the mutation site (amino acid residues 69-76). Structural analysis of the MA trimer revealed that the side chain of Thr^70^, which resides in the loop connecting helices 4 and 5, is sandwiched between the side chains of Gln^59^ and Gln^63^ of a neighboring molecule (Fig. 4). Interestingly, the amide resonances of these two residues did not exhibit significant chemical shift changes. On the other hand, mutation of Leu^75^ to Gly induced substantial CSPs that extended beyond the mutation site, including amino acid residues in the trimer interface (Figs. 3 and 4). Of note, residue Leu^75^, located in helix 5, is not involved in intermolecular interactions with a neighboring molecule. However, within the same molecule the side chain of Leu^75^ is relatively close to the side chain of Leu^64^ located in helix 5 (4 Å) and is spatially close to the loop made of residues 67-72. As discussed above, amino residues 67-72 are located in the trimer interface and are in close proximity to the loop made of amino acid residues 43-47. Therefore, it is conceivable that Leu^75^ contributes to the orientation of the loop.

Taken together, NMR data provided insights into structural perturbations within the trimer interface. However, we were unable to precisely determine the structural changes in the trimer interface by NMR methods because, like the WT MA protein (26), detection of intermolecular contacts (e.g. NOE) proved to be technically challenging. In summary, we provided structural evidence that substitution of amino acid residues in the trimer interface can directly or indirectly contribute to the (de)stabilization of the MA trimer form.

### Crystal structure of the myr(–)MA Q63R protein

As discussed above, to explain the role of MA trimerization in Env incorporation, it was suggested that the Q63R mutation in MA may stabilize the trimer structure such that MA lattices which form large hexamer holes are favored over those that feature small hexamer holes (Fig. 1) (44). However, it is not clear how this substitution can stabilize the trimer form. We have shown that, similar to the WT MA protein, MA Q63R is in monomer–trimer equilibrium. As indicated by the NMR data, substitution of Gln^63^ to Arg did not induce significant CSPs of residues in the trimeric interface. To be able to assess the structural changes in the MA trimer, we determined the high-resolution structure of myr(–)MA Q63R protein by x-ray crystallography. Atomic coordinates and structure factors for two crystal forms have been deposited in the PDB (codes 7JXR and 7JXS; Table 1). Interestingly, the two crystal forms of myr(–)MA Q63R yielded structures that are essentially identical with slightly different parameters (Fig. S2 and Table 1). Each asymmetric unit of the crystal structure contains two trimers. The structure of the myr(–)MA Q63R trimer in both crystal forms is virtually identical to that of the WT myr(–)MA protein with slight orientation differences of monomers in the trimeric assembly (Fig. 5) (21). The structures in the two crystal forms revealed only minor conformational changes in the loops connecting helices I and II and helices V and VI (Fig. S2). Intriguingly, the guanidinium group of Arg^63^ is 2.54 Å (averaged across 12 examples in the asymmetric unit of the two crystal forms) from the hydroxyl group of Ser^67^ of an adjacent MA molecule, which forms new hydrogen bonding in the trimer interface (Fig. 5). This result demonstrates that substitution of Gln^63^ with Arg stabilizes the trimer structure of MA through H-bonding between the side chains of Arg^63^ and Ser^67^ located on two neighboring MA molecules.

**Figure 5.**
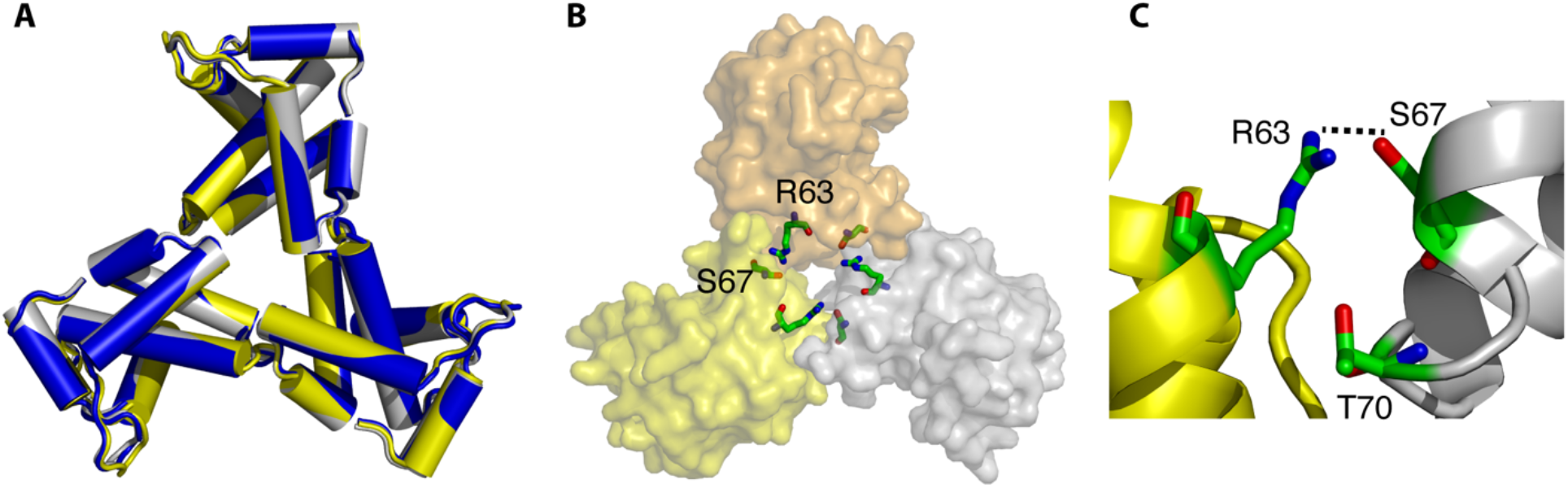
High-resolution crystal structure of the HIV-1 myr(–)MA Q63R protein. *A*, superposition of WT myr(–)MA trimer protein (PDB code 1HIW; blue) and the two trimers found in the asymmetric unit of the crystal structure of myr(–)MA Q63R (yellow and grey). The structure of the myr(–)MA Q63R mutant is essentially identical to that of the WT myr(–)MA protein. *B*, surface representation of the trimer arrangement of the myr(–)MA Q63R protein. Amino acid residues Arg^63^ and Ser^67^ are shown as sticks. *C*, cartoon representation of myr(–)MA Q63R mutant showing the H-hydrogen bond formed between the side chains of Arg^63^ and Ser^67^. Residue Thr^70^ is also shown to highlight the close proximity to Arg^63^.

**Table 1.** X-ray diffraction and refinement statistics

| **Parameter** | **Crystal 1** | **Crystal 2** |
| --- | --- | --- |
| Wavelength | 1.000 | 1.000 |
| Resolution range | 46.21–2.04 (2.113–2.04) | 36.82–2.35 (2.434–2.35) |
| Space group | P 1 21 1 | P 1 21 1 |
| Unit cell | 63.13 97.92 84.81 90 99.94 90 | 64.54 90.72 73.16 90 102.38 90 |
| Total reflections | 171742 | 100393 |
| Unique reflections | 61490 (6078) | 32573 (3286) |
| Multiplicity | 2.8 | 3.1 |
| Completeness (%) | 95.03 (94.19) | 94.51 (95.36) |
| Mean I/sigma(I) | 28.86 (2.50) | 25.95 (2.38) |
| Wilson B-factor | 42.84 | 49.51 |
| R-merge | 0.067 (0.408) | 0.072 (0.431) |
| R-meas | 0.083 (0.508) | 0.088 (0.515) |
| R-pim | 0.048 (0.298) | 0.050 (0.277) |
| CC1/2 | 0.987 (0.726) | 0.987 (0.807) |
| Reflections used in refinement | 61481 (6078) | 32566 (3285) |
| Reflections used for R-free | 1995 (197) | 1999 (202) |
| R-work | 0.2032 (0.2525) | 0.2170 (0.2481) |
| R-free | 0.2340 (0.3166) | 0.2560 (0.2723) |
| Number of non-hydrogen atoms | 5507 | 5286 |
| macromolecules | 5141 | 5111 |
| ligands | 149 | 108 |
| solvent | 217 | 67 |
| Protein residues | 638 | 632 |
| RMS(bonds) | 0.008 | 0.008 |
| RMS(angles) | 1.2 | 1.07 |
| Ramachandran favored (%) | 100 | 98.06 |
| Ramachandran allowed (%) | 0.00 | 1.61 |
| Ramachandran outliers (%) | 0.00 | 0.32 |
| Rotamer outliers (%) | 1.07 | 3.22 |
| Clashscore | 5.99 | 8.92 |
| Average B-factor | 51.98 | 65.77 |
| macromolecules | 51.67 | 65.83 |
| ligands | 59.99 | 65.32 |
| solvent | 53.77 | 61.45 |
| Number of TLS groups | 40 | 29 |

The values in parentheses correspond to the highest resolution data shell.

### Oligomeric properties of MA mutants

As mentioned earlier, previous sedimentation equilibrium (SE) studies have shown that, while myr(–)MA is monomeric at all tested protein concentrations and pH values (22–27), the MA protein is in monomer–trimer equilibrium (24, 26), a process that is modulated by changing solution pH (27). These results were explained by a (de)protonation process of the His^89^ imidazole ring. Deprotonation of His^89^ destabilizes the salt bridge formed between His^89^(Hδ2) and Glu^12^(COO-), leading to tight sequestration of the myr group and a shift in the equilibrium from trimer to monomer. To determine how the various MA mutations impacted the oligomerization properties of the MA protein in solution, we collected sedimentation velocity (SV) and SE data on the WT and mutant MA proteins at pH 5.5. Consistent with previous data, the SV profile for myr(–)MA exhibits a sharp sedimentation boundary indicating a monomeric character (Fig. 6). The MA protein, however, exhibits a broad sedimentation boundary with a larger S value consistent with a monomer–trimer equilibrium. The MA mutants exhibited slightly variable sedimentation boundaries with an overall sedimentation coefficient (c(s)) distribution similar to that of the WT MA protein. Interestingly, the sedimentation boundary for the L75G mutant is sharper than the WT and the other mutants and shifted to a smaller S value, suggesting that the monomer–trimer equilibrium is shifted towards the monomeric form (Fig. 6). Next, we collected SE data for WT and mutant MA proteins at pH 5.5. Like the WT protein, SE data for all mutants best fit a monomer–trimer model. Interestingly, while the *K*_a_ values for the A45E, T70R and Q63R mutant proteins are similar to the WT protein, the corresponding value for the L75G mutant is ~3-fold lower (Fig. 6), consistent with a lower trimer population. Taken together, the SV and SE data indicate that whereas none of the four mutations in MA significantly altered the monomer–trimer equilibrium in solution, the L75G mutation appears to shift the equilibrium towards the monomer form.

**Figure 6.**
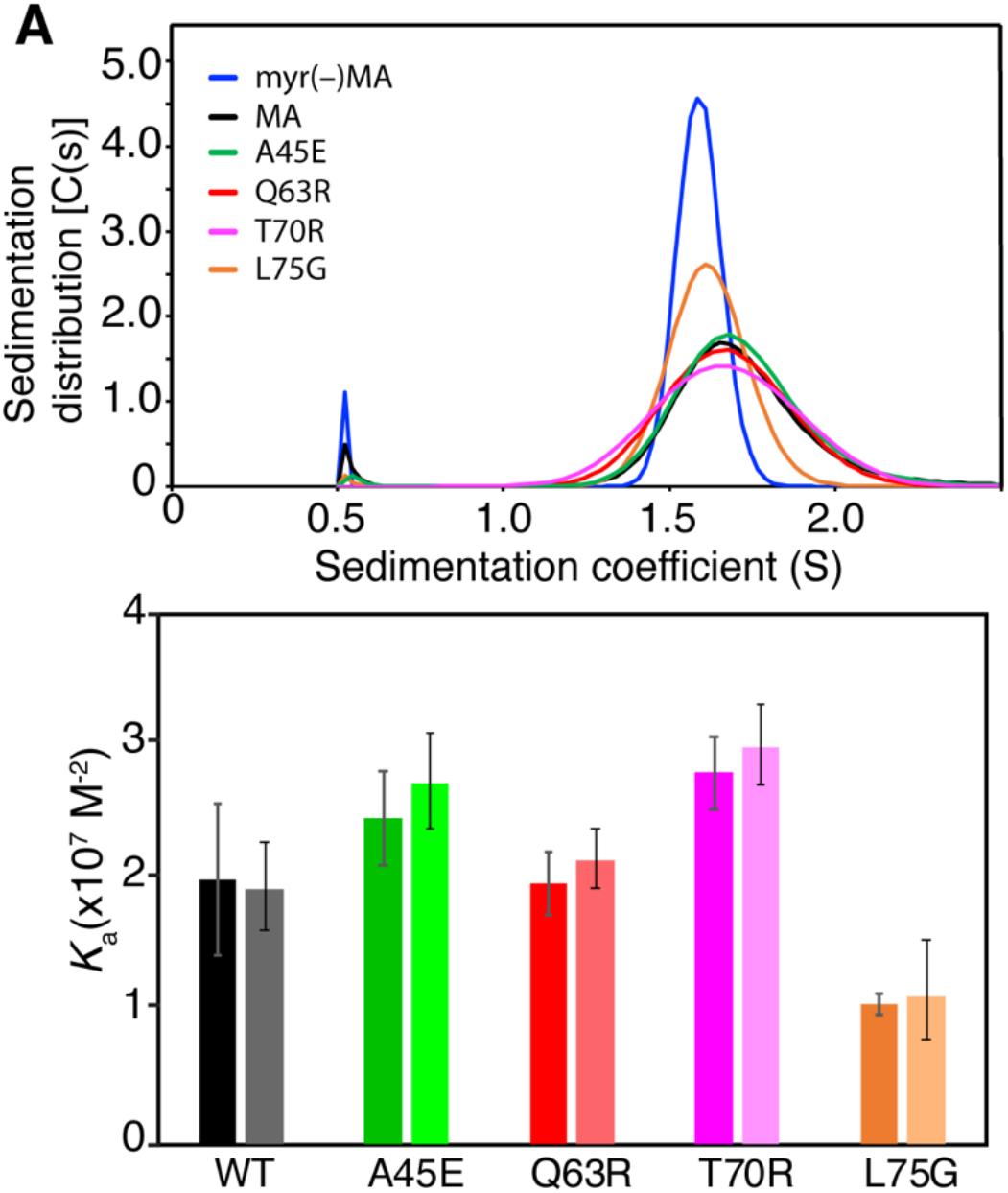
Sedimentation velocity and equilibrium data. *A*, Sedimentation coefficient distributions, c(s), obtained from the sedimentation profiles for WT and mutant MA at pH 5.5. *B*, A histogram showing the monomer–trimer association constants obtained from sedimentation equilibrium experiments at pH 5.5. The left series (dark shades) show the average *K*_a_ and standard deviation from 2-3 replicates of the SE experiment, and the right series (light shades) show the *K*_a_ for a single SE fit which had the closest value to the mean *K*_a_ for replicates, along with the F-statistic confidence intervals at 95.46% for the fitted value of *K*_a_.

## Discussion

Point mutations of MA residues Leu^13^, Glu^17^, Leu^31^, Val^35^, and Glu^99^, located at the tip of the hexamer centers in the hexamer of trimer model (Fig. 1), were found to impair Env incorporation without affecting virus particle formation (54–57). Freed and co-workers have provided biochemical evidence that MA trimerization is an obligatory step in the assembly of infectious HIV-1 and demonstrated a correlation between loss of MA trimerization and loss of Env incorporation (44). Nonconservative mutations at, or near, the trimer interface in the crystal structure (Fig. 1) inhibited MA trimerization and yielded particles with impaired Env incorporation and infectivity (44). Substitution of residue Gln^63^ with Arg suppressed Env incorporation defects of the L13E, E17K, L31E, V35E, and E99V MA mutations (42–44, 50). The current hypothesis suggests that Q63R mutation may stabilize the trimer structure such that MA lattices, which form large hexamer holes, are favored over those that feature small hexamer holes (Fig. 1) (44). This hypothesis was further supported by the finding that Q63R mutation promoted interaction with gp41CT without altering the organization of MA on a membrane layer (41). A correlation between MA trimerization and gp41CT binding was also suggested in a biochemical study involving MA mutants and MACA proteins, supplemented with inositol polyphosphates (64).

In this study, we have shown that: (i) mutations that either impaired (A45E, T70R, and L75G) or rescued (Q63R) Env incorporation had no adverse effects on the structures and folding of the MA protein. This result rules out the possibility that such mutations produce a nonfunctional protein that indirectly impacts its ability to form a trimer or participate in other functions involved in Env incorporation. (ii) None of the mutations had any detectable effect on the efficiency of myristoylation or concentration–dependent myr switch. NMR data obtained at different protein concentrations indicate that all four mutants display a concentration–dependent myr switch mechanism, similar to the WT MA protein. (iii) Mutations appear to induce minor structural perturbations in the trimer interface, which can directly or indirectly contribute to the (de)stabilization of the MA trimer form. As detected by NMR, the largest CSPs were observed for the L75G mutant. This result is not surprising given that Leu^75^ is located in a helical motif. Substitution with a glycine reside, which is sometimes known as a helix breaker, may have induced a larger structural change than the other three mutants. As mentioned earlier, the effect of Leu^75^ mutation on MA trimerization is probably indirect as this residue is not involved in intermolecular interactions with a neighboring MA molecule. Leu^75^ is in close proximity to Leu^64^, which is in juxtaposition to the loop made of amino acid residues 67-72 that interacts with amino acid residues 42-46 of a neighboring molecule. Therefore, it is conceivable that Leu^75^ contributes to the orientation of this loop to enable interaction with an adjacent molecule. (iv) A new hydrogen bond is formed between the side chains of Arg^63^ and Ser^67^ in the center of the trimer interface of myr(–)MA Q63R, providing the first structural evidence for a stabilization of the trimer form. (v) SV and SE data show that the A45E, T70R, and Q63R mutations had minimal effect on the monomer–trimer equilibrium, whereas the L75G mutation enhanced the population of the monomeric form.

As discussed above, the structures of MA and myr(–)MA are nearly identical (21, 22, 25, 26, 65). The only distinct difference is that the X-ray structure of myr(–)MA revealed a trimer arrangement of MA molecules (21). The MA protein, however, is in monomer–trimer equilibrium. To determine whether the MA–MA interface in solution resembles that in the x-ray structure of myr(–)MA (21), we recently engineered a stable MA trimer by fusing a FD domain on the C-terminus of MA (28). Hydrogen–deuterium exchange MS data supported a MA–MA interface that is consistent with that observed in the crystal structure of the myr(–)MA trimer (28). Of note, the region exhibited a significant increase in protection from deuteration in the HDX-MS assay is the same region identified here as a putative trimer interface. Our data presented here demonstrate that neither of the three mutants that were proposed to inhibit MA trimerization (A45E, T70R, and L75G) completely shifted the equilibrium to a monomer, nor did the Q63R mutation which rescues Env incorporation, forms a stable trimer in solution even though a new H-bond is observed between the side chains of Arg^63^ and Ser^67^ from neighboring subunits (Fig. 5). This discrepancy between the crystal structure and solution data is still not well understood.

It is important to highlight the link between the structural findings and the previous biological findings. It is worth noting that previous studies indicated that residue Gln^63^ is not crucial for Env incorporation in the context of otherwise-WT MA, and that Q63R is unique for its ability to fully rescue Env incorporation defects (42). It appears that the type of amino acid substitution at position Gln^63^ is key for its function in restoring Env incorporation. When introduced as a single mutation in MA, Q63R only minimally enhanced Env incorporation (42). However, the effect was more dramatic in restoring Env incorporation to a WT level when this mutation was introduced along with the Env-defective L13E mutant (42). By substituting Gln^63^ with E, G, K, L, N, or W, it was shown that all mutants replicated with WT kinetics in Jurkat cells, and that none of the single mutations severely impaired virus release, infectivity or Env incorporation. When the L13E/Q63[E/G/K/L/N/W] double mutants were subjected to the same analysis, none of the Gln^63^ mutants was able to fully rescue the virus replication, infectivity, and Env incorporation defects imposed by the L13E mutant (42). Interestingly, even the L13E/Q63K mutant exhibited a partial rescue, as infectivity in TZM-bl cells was comparable to that of L13E/ Q63R, no rescue of Env incorporation was apparent. Taken together, Q63R appears to be unique for its ability to fully rescue Env incorporation defects. Our findings complemented the biological data by showing that the H-bond formed between the side chains of Arg^63^ and Ser^67^ is key for the stability of the MA trimer and hence to its function in rescuing Env incorporation (42). It is possible that slight changes to trimerization affinity may be amplified in a biological context, due to the 2D constraint imposed by the membrane or the presence of other oligomerization promoting domains of Gag such as CA.

Role of Ser^67^ and Thr^70^ in Env incorporation in the context of WT, L13E, Q63R, and L13E/Q63R clones was also investigated (42). Whereas the S67A mutation had no effect on the phenotypes of the four viral clones, T70A blocked the ability of Q63R to rescue L13E infectivity and Env incorporation, although Q63R/T70A was as infectious as Q63R alone. The single T70A mutant was also impaired for infectivity and Env incorporation, suggesting that Thr^70^ may be involved in MA function. Another interesting result is the finding that S67R behaved like Q63R in its ability to rescue the defect imposed by L13E, indicating that an Arg at either residue 63 or 67 could rescue the L13E defect (42). Based on the structure of the myr(–)MA Q63R protein (Fig. 5), it is clear that Ser^67^ plays an important role in forming a H-bond with Arg^63^. It is expected that an Arg at position 67 is also capable of forming a H-bond with the side chain of Gln^63^ on a neighboring molecule, therefore stabilizing the MA trimer structure. The close proximity of Arg^63^ and Ser^67^ and their role in the stabilization of the MA trimer is supported by the finding that MA forms dimer and trimer in the glutaraldehyde cross-linking assay conducted with the double mutant (Q63K/S67K) in Jurkat and MT4 cell lines (44).

Combining previous studies with the data presented here, it appears that the correlation between trimerization of the MA domain of Gag and Env incorporation is not straightforward. Mutation of residues located in the trimer interface, on the tip of the hexamer ring, or even distant residues can impact Env incorporation. In a recent study, new compensatory mutations that rescue MA trimer interface mutants with severely impaired Env incorporation in MT-4 T-cells were identified (45). Viruses with MA L75G mutation acquired mutations G11S, V35I/F44I, V35I/E52K, V46I and E52K. For viruses with MA L75E mutation, compensatory mutations D14N/V45I, Q28K, V35I/F44L and E52K were identified. Of note, among all these secondary mutations only Phe^44^ and Val^46^ are located in the trimer interface. It was suggested that mutations that are distant from the trimer interface may indirectly influence the trimer interface by inducing potential structural or conformational changes (45). Structural studies to discern conformational changes induced by the compensatory mutations are warranted.

Our data support the hypothesis that mutations in MA that block Env incorporation do so by disrupting an otherwise-stable MA lattice, rather than by disrupting a specific MA–gp41 interaction. The importance of the trimer interface in rescuing Env incorporation suggests that a stable MA trimer is key to trap Env into the Gag lattice. This model is also supported by recent high-resolution single-molecule tracking data (66), which demonstrated that Env trimers are confined to subviral regions of a budding Gag lattice, supporting a model where direct interactions and/or steric corralling between the gp41CT and MA lattice promote Env trapping. In summary, the new findings advance our understanding of the complementary, yet distinct, roles the MA domain of Gag and gp41CT play in Env incorporation.

## Experimental procedures

### Plasmid Construction and Protein Expression

A plasmid encoding for HIV-1 MA (pNL4-3 strain) and yeast N-terminal myristoyl transferase was provided by Dr. Michael Summers (Howard Hughes Medical Institute, University of Maryland Baltimore County, MD). The A45E, T70R, Q63R and L75G MA mutant constructs were generated using a QuickChange XL site-directed mutagenesis kit (Stratagene). Forward and reverse primers (Integrated DNA Technologies) extended 15 base-pairs on either side of the mutation codon. Mutations were verified by plasmid sequencing at the Heflin Genomics Core at the University of Alabama at Birmingham. WT and mutant MA, and myr(–)MA proteins were prepared as described (22, 24–26).

### NMR Spectroscopy

NMR data were collected at 35 °C on a Bruker Avance II (700 MHz ^1^H) or Avance III (600 or 850 MHz ^1^H) spectrometers equipped with cryogenic triple-resonance probes, processed with NMRPIPE (67) and analyzed with NMRVIEW (68) or CCPN Analysis (69). The backbone resonances were assigned using standard triple resonance data (HNCA, HN(CO)CA, HN(CO)CACB, and HNCACB) collected at 35 °C on 100–500 μM samples in a buffer containing 50 mM sodium phosphates (pH 5.5), 50 mM NaCl, and 1 mM tris(2-carboxyethyl)phosphine hydrochloride (TCEP). The triple-resonance experiments were collected as non-uniformly sampled (NUS) sparse data (20% sampling density in indirect dimensions) according to schemes generated using hmsIST (70). Chemical shift perturbations were calculated as 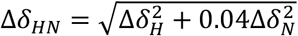, where Δ*δ_H_* and Δ*δ_N_* are ^1^H and ^15^N chemical shift changes, respectively. Histograms for the chemical shift changes were generated using gnuplot software (http://www.gnuplot.info) (71). Structural representations of the chemical shift changes were generated using PyMOL (Version 2.3.2 Schrödinger, LLC) (72).

### X-ray crystallography and structure determination

The HIV-1 myr(–)MA Q63R protein used in the crystallization trials was at 20 mg/mL in a buffer containing 10 mM Tris.HCl (pH 8), 1 mM EDTA, 2 mM β-mercaptoethanol, and 50 mM NaCl. Diffraction quality crystals were obtained using hanging-drop vapor diffusion in a solution of 40% PEG (molecular weight 400), 0.1 M sodium acetate (pH 4), and 50 mM lithium sulfate. Crystals were cryo-cooled in the same conditions. X-ray diffraction data were collected at the Advance Photon Source, SER-Cat Beamline 22-ID. Raw intensity data were processed with the HKL2000 software package (73). The initial electron density map was generated via molecular replacement with PHASER (74) using the previously solved structure of the WT myr(–)MA protein (PDB code 1HIW). The structure was then iteratively refined with PHENIX (75) and Coot (76). Visualization of structures was performed using Pymol.

### Analytical Ultracentrifugation

Sedimentation velocity (SV) and sedimentation equilibrium (SE) experiments were performed on a Beckman XL-I Optima ultracentrifuge equipped with a four-hole An-60 rotor (Beckman Coulter, Pasadena, CA). Cells were equipped with double-sector, charcoal-filled epon centerpieces with 12 mm path lengths and sapphire windows. Prior to AUC data collection, protein samples were run on a size exclusion chromatography column (Superdex 75, 10/300 GL, cytiva) in a buffer containing 50 mM sodium phosphates (pH 5.5), 100 mM NaCl and 2 mM TCEP. Loading concentrations of MA samples ranged from 30–100 μM. For SV experiments, rotor speed was set at 40,000 rpm. SE experiments were performed at 22000, 26000, and 30000 rpm. All experiments were carried out at 20 °C. Scans were collected at a wavelength of 280 nm for SV and 250 nm for SE. Partial specific volumes (v-bar) and molar extinction coefficients were calculated by using the program SENDTERP, and buffer densities were measured pycnometrically. SV data were analyzed using SEDFIT (77–80), and SE data were analyzed with HETEROANALYSIS (81). Sedimentation coefficients obtained from SV experiments were corrected to 20 °C and infinite dilution in water (*s_20,w_*). Equilibrium association constants were obtained by global fitting of SE scans collected at a single concentration and at three rotor speeds using a monomer–trimer equilibrium model. The molecular weight of the monomer (15.6 kDa) and the complex stoichiometry (n = 3) were treated as fixed parameters. Reference concentrations and baselines for each scan were floated parameters, and the *K*_a_ for the monomer–trimer equilibrium was fit as a global parameter. The confidence intervals for the *K*_a_ at 95% were determined using the F-statistic method in HETEROANALYSIS (81).

## Acknowledgments

We thank Dr. Aaron Lucius (University of Alabama at Birmingham) for helping with the AUC experiments. We also the O’Neal Comprehensive Cancer Center at the University of Alabama at Birmingham (funded by the NCI grant P30 CA013148) for supporting the High-Field NMR facility. X-ray data were collected at the Southeast Regional Collaborative Access Team (SER-CAT) 22-ID beamline at the Advanced Photon Source, Argonne National Laboratory. SER-CAT is supported by its member institutions, and equipment grants (S10_RR25528 and S10_RR028976) from the National Institutes of Health. Use of the Advanced Photon Source was supported by the U. S. Department of Energy, Office of Science, Office of Basic Energy Sciences, under Contract No. W-31-109-Eng-38.

## Author contributions

GNE, RHG, TJG and JSS designed the experiments. GNE and RHG expressed, purified, and characterized the proteins. GNE and JSS performed the NMR experiments and analyzed the results. TJG collected and analyzed the x-ray data. GNE and JSS wrote the paper. GNE, TJG and JSS edited the paper.

## Footnotes

This work was supported by grants 5R01GM117837 and 9R01AI150901 from the National Institutes of Health (NIH) to JSS. The High-Field NMR facility at the University of Alabama at Birmingham was established through the NIH (1S10RR026478) and is currently supported by the O’Neal comprehensive cancer center (NCI grant P30 CA013148).

The atomic coordinates and structure factors (codes 7JXR and 7JXS) have been deposited in the Protein Data Bank (http://wwpdb.org/).

## The abbreviations used are

MA: myristoylated matrix
myr(–)MA: unmyristoylated matrix
PI(4,5)P_2_: phosphatidylinositol 4,5-bisphosphate
NMR: nuclear magnetic resonance
HSQC: heteronuclear single quantum coherence
CSP: chemical shift perturbation
AUC: analytical ultracentrifugation

## Supporting Information

**Fig. S1.**
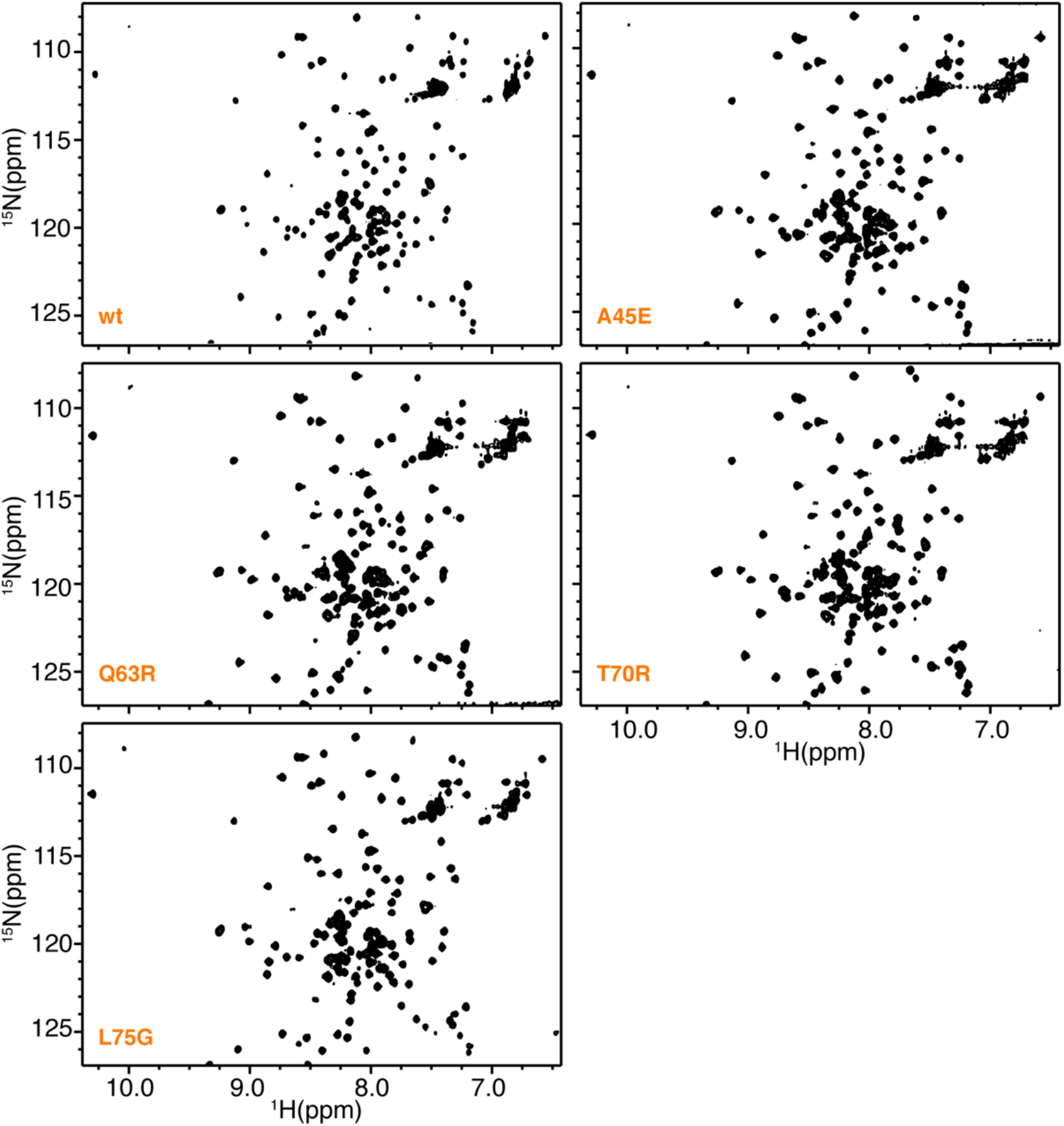
2D ^1^H-^15^N HSQC spectra obtained for WT and mutant HIV-1 MA proteins at 150 μM (308 K) in a buffer containing 50 mM sodium phosphates (pH 6) and 50 mM NaCl.

**Fig. S2.**
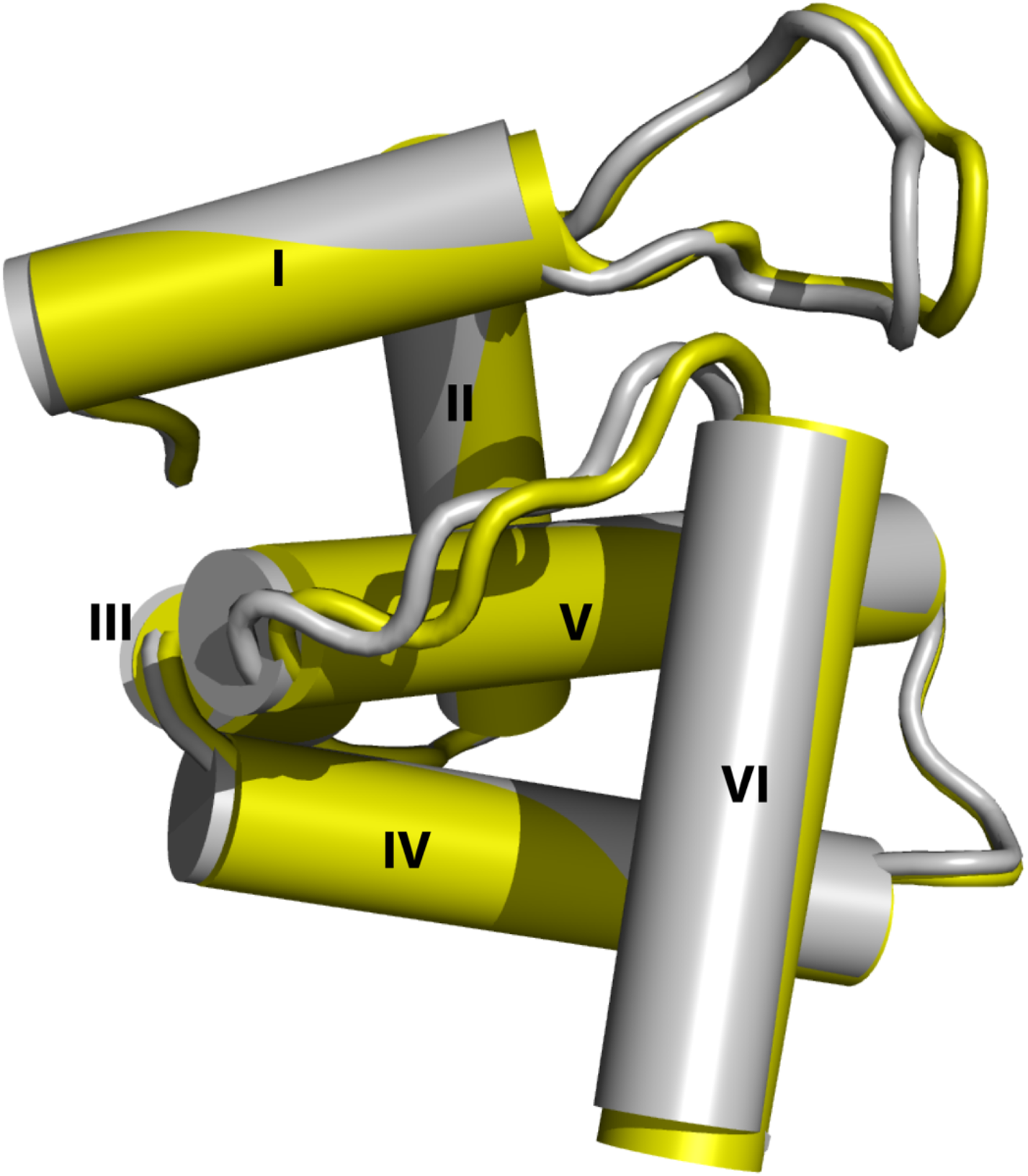
Superimposition of HIV-1 myr(–)MA Q63R structures from crystals form 1 (PDB 7JXR; yellow) and form 2 (PDB 7JXS; grey). Residues 110-132 are not shown. Ribbon representations of the structures were generated using the PyMOL molecular graphics system (Schrödinger, LLC).

